# A non-adaptive explanation for macroevolutionary patterns in the evolution of complex multicellularity

**DOI:** 10.1101/2023.11.11.566713

**Authors:** Emma P. Bingham, William C. Ratcliff

## Abstract

“Complex multicellularity”, conventionally defined as large organisms with many specialized cell types, has evolved five times independently in eukaryotes, but never within prokaryotes. A number hypotheses have been proposed to explain this phenomenon, most of which posit that eukaryotes evolved key traits (*e*.*g*., dynamic cytoskeletons, alternative mechanisms of gene regulation, or subcellular compartments) which were a necessary prerequisite for the evolution of complex multicellularity. Here we propose an alternative, non-adaptive hypothesis for this broad macroevolutionary pattern. By binning cells into groups with finite genetic bottlenecks between generations, the evolution of multicellularity greatly reduces the effective population size (*Ne*) of cellular populations, increasing the role of genetic drift in evolutionary change. While both prokaryotes and eukaryotes experience this phenomenon, they have opposite responses to drift: mutational biases in eukaryotes tend to drive genomic expansion, providing additional raw genetic material for subsequent multicellular innovation, while prokaryotes generally face genomic erosion. These effects become more severe as organisms evolve larger size and more stringent genetic bottlenecks between generations— both of which are hallmarks of complex multicellularity. Taken together, we hypothesize that it is these idiosyncratic lineagespecific mutational biases, rather than cell-biological innovations within eukaryotes, that underpins the long-term divergent evolution of complex multicellularity across the tree of life.

## Introduction

Multicellularity has evolved over 50 times independently, arising at least 3 billion years ago in cyanobacteria (1). Most of these lineages have remained relatively simple, with “complex” multicellularity arising only five times, all within eukaryotes (animals, plants, fungi, brown, and red algae). The most ancient of these transitions are the red algae, which began evolving multicellularity ∼1.1 billion years ago (2). Complex multicellularity is strikingly absent in the other two domains of life, bacteria and archaea, despite their 2-billion-year head start.

The dramatic difference between multicellular prokaryotes and eukaryotes is widely acknowledged, and begs an evolutionary explanation. Many hypotheses have been proposed to explain the success of multicellularity within eukaryotes, most of which highlight specific cell biological innovations within this domain. For example, eukaryotes have a more dynamic cytoskeleton, which may facilitate differentiation and adhesion (3). Introns permit alternative splicing of transcripts, increasing the potential number of functional proteins encoded by a genome (4). Eukaryotes evolved additional mechanisms of gene regulation, such as miRNAs, which allow precise control of gene expression and likely facilitated cellular differentiation during multicellular evolution (3). Compartmentalization within organelles partition metabolic and gene regulatory space, allowing novel functions to evolve (5, 6). Prior work has argued that the evolution of mitochondria fueled genome expansion by increasing energy availability (7), though more recent work argues that this conclusion is based on faulty assumptions (8).

While there is little doubt that eukaryotic traits listed above have played a role in the evolution of complex multicellularity, it is not clear that they are a firm requirement. Prokaryotes are diverse, and as a group possess many functional cell-biological tools that could be exapted for novel multicellular morphogenesis, including mechanisms for cell-cell communication, differentiation, and collective growth (9). So why, then, haven’t multicellular prokaryotes evolved complex multicellularity?

Here, we propose that a key factor underlying the repeated evolution of complex multicellularity in eukaryotes, and its absence in prokaryotes, stems from their divergent genomic responses to reduced effective population size (*Ne*). When multicellular groups evolve from unicellular ancestors, *Ne* decreases dramatically, due to reproductive bottlenecks as cells are partitioned into groups. This is especially pronounced in clonal multicellular organisms with stringent genetic bottlenecks between generations. This reduction of *Ne* has divergent consequences in eukaryotes compared to prokaryotes: mutational biases within eukaryotes lead to genomic expansion under low *Ne*, while prokaryotes are biased towards gene deletion and genomic erosion. This fundamental dichotomy may facilitate the evolution of complex multicellularity in eukaryotes, while inhibiting it in prokaryotes.

## Results and Discussion

### Clonal Multicellularity Reduces Ne

All extant lineages of complex multicellularity develop clonally, rather than through aggregation of unrelated cells. This is well-explained by evolutionary theory as a key trait facilitating multicellular adaptation and cellular specialization (10). Clonal multicellularity, however, greatly reduces *Ne*. Unicellular populations contain large numbers of independently-reproducing individuals. Once arranged into multicellular groups, cells are partitioned into discrete units that undergo bottlenecks during organismal reproduction. The extent to which *Ne* is reduced by multicellularity depends both on organismal size and the severity of the reproductive bottlenecks between generations.

While precise estimates of *Ne* depend on specific details of population structure and dynamics, we can examine the impact of multicellularity on the capacity for a population to support genetically distinct lineages, a key component of *Ne*, with a simple model. Specifically, if we assume that the total number of cells in a population is fixed (*i*.*e*., carrying capacity scales with biomass, and thus the number of cells, not the number of groups they are in), then we can calculate the maximum number of genetically distinct lineages that a population of *N* cells can support, *L*, as:

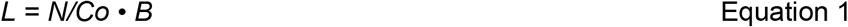

where *Co* is the number of cells in each organism, and *B* is the bottleneck size (number of cells transferred to the next generation) at reproduction.

The evolution of large organisms that undergo small genetic bottlenecks during reproduction radically reduces the potential genetic diversity of populations (Figure 1a). Consistent with this theory, empirical measurements of *Ne* show that it is typically 2-3 orders of magnitude lower in multicellular populations, relative to unicellular taxa (11). Thus the evolution of multicellularity, and especially the evolution of macroscopic organismal size and unicellular genetic bottlenecks during ontogeny, radically changes the population genetic context in which evolution occurs.

**Figure 1.**
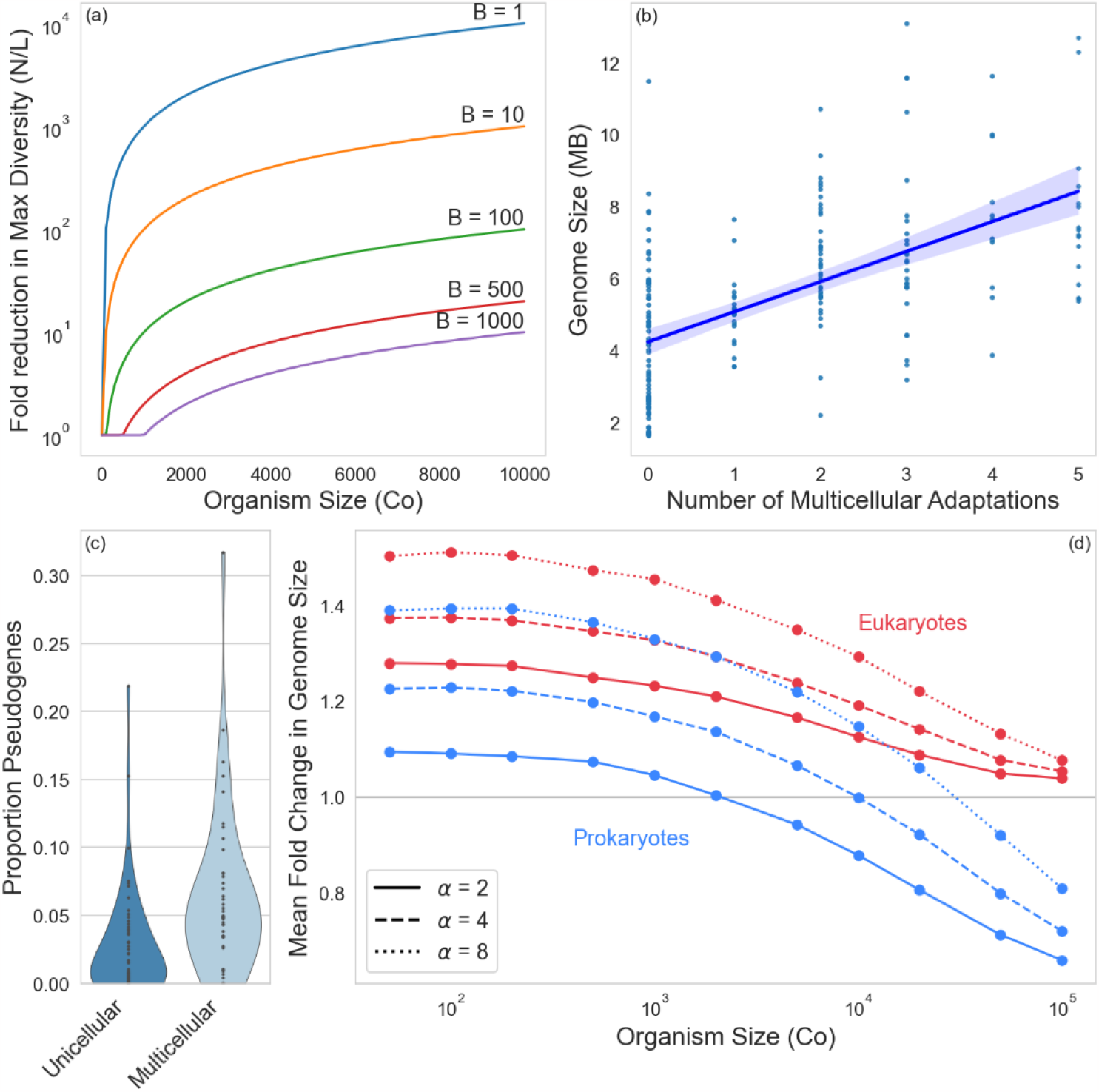
Population genetic consequences of multicellular evolution. (a) The evolution of clonal multicellularity greatly reduces the effective population size of cellular populations, shown here as the difference between the census cellular population size *N* and the maximum number of genetically-distinct lineages, *L* (Equation 1) in the population. Different lines reflect different bottleneck sizes, *B*. (b) Comparative genomics and evolutionary theory predict that, all else equal, the evolution of increasingly complex multicellularity requires genomic expansion via the evolution of additional protein coding genes and regulatory elements. Indeed, cyanobacteria with a larger number of multicellular adaptations (see Methods) have larger genomes (shown is an ordinary least squares regression with a shaded 95% confidence interval). (c) Multicellular cyanobacteria, however, had more than twice the frequency of pseudogenes than unicellular taxa, consistent with the hypothesis that in bacteria, multicellularity-driven reductions in *Ne* lead to genome degradation. (d) The combination of multicellularity-driven reduced *Ne* and a deletion bias in prokaryotes can prevent even strong selection from driving increased genome size. Using a Wright-Fisher simulation, we considered a scenario where larger genome size is adaptive in nascent multicellular organisms (*α* is a scaling factor modifying the positive interaction between genome size and organism size, and in this simulation, *B* = 1, see Methods). Increased genetic drift from a multicellular life history prevents genomic expansion in larger organisms, especially prokaryotes, despite strong selection for greater genome size.

### Prokaryotes and Eukaryotes Show Opposite Genomic Responses to Low Ne

While both prokaryotes and eukaryotes experience greater drift with lower *Ne*, they exhibit fundamentally different genomic consequences as a result. Eukaryotes tend to expand their genomes, while prokaryotic genomes tend to shrink (12).

20 years ago, Lynch and Conery (2003) argued in a groundbreaking paper that genetic drift plays a fundamental role in the evolution of larger genome size in eukaryotes (13). In eukaryotes, weakened efficacy of selection allows proliferation of mobile genetic elements like transposons. Purifying selection also becomes less effective at removing introns, leading to intron bloat. And gene duplicates are more likely to reach fixation when ineffective purifying selection permits subfunctionalization or neofunctionalization.

Recently, it has become clear that prokaryotes have the opposite response to genetic drift: they generally have an innate bias towards eliminating portions of their genome through deletions (14). Prokaryotes experience high rates of replication fork collapse leading to slippage-mediated sequence loss. Many species lack non-homologous end joining, which can cause insertions in eukaryotes. Together with recombination between repetitive sequences that tends to cause deletions, the major mutational mechanisms in prokaryotes favor the removal of DNA. This innate deletion bias, coupled with ineffective selection against gene loss under low *Ne*, drives the incremental erosion of prokaryotic genomes when *Ne* is small.

### Consequences for Multicellularity

This fundamental difference in how prokaryotic and eukaryotic genomes respond to reduced *Ne* has fundamental consequences for lineages transitioning to multicellularity. In nascent multicellular eukaryotes, the lowered *Ne* imposed by reproductive bottlenecks would have driven expansions of genomic content. This drift-induced increase in genetic material could provide fodder for evolutionary innovation through emergence of new genes, regulatory elements, and splice variants (13).

Conversely, in prokaryotes, the evolution of multicellularity initiates an uphill battle against degradative evolution driven by nonadaptive genome streamlining. While selection for increased multicellular functionality would have favored larger genomes, the concomitant reduction in *Ne* would erode genomic content through pseudogenization. Even more problematically, this process would have only been exacerbated by the evolution of larger organismal size and stringent genetic bottlenecks (Figure 1a)— both hallmarks of complex multicellularity.

The evolution of increasingly complex multicellularity is usually associated with a substantial expansion of genome size and content. In fungi, which are a useful lineage for this comparison as multicellularity has been lost secondarily in multiple clades, taxa with complex multicellularity typically have more than twice as many protein coding genes as unicellular relatives (15). Similarly, plants, animals and macroalgae have substantially larger genomes than their unicellular relatives (11, 13), both in terms of the number of protein coding genes (16, 17) and non-coding regions, which serve as a crucial substrate for the evolution of regulatory regions underpinning cellular specialization and morphogenesis (18).

To explore the tension between adaptive genome expansion in multicellular bacteria, and non-adaptive genome degradation that arises as a side-effect of increased genetic drift, we examined the oldest, and arguably most paradigmatic lineage of prokaryotic multicellularity: cyanobacteria. Prior work has identified key steps in the evolution of multicellularity in this clade (1), identifying five multicellular adaptations: filamentous growth (multicellularity *sensu stricto*), gas vesicles, true branching, akinetes and hormogonia. Consistent with expectations, we see a strong positive correlation (*r* = 0.57) between the number of multicellular adaptations an organism possesses and its overall genome size (Figure 1b, *y* = 8.3·10^5^*x* + 4.2·10^6^, *p* = 5·10^-19^, ordinary least squares regression), underscoring the importance of genome expansion for multicellular adaptation. Multicellular cyanobacteria, however, had a 2.3-fold greater proportion of pseudogenes than unicellular species (*t*_*98*_ = 3.4, *p* = 0.001, two sample *t*-test; Figure 1c), consistent with the hypothesis that the evolution of multicellularity, and the corresponding reduction in *Ne*, is associated with genome degradation in bacteria.

The above results suggest that the deletional bias of prokaryotic genomes powerfully constrains the joint evolution of large multicellular organisms with large genomes size-both of which are hallmarks complex multicellularity. To further test this hypothesis, we examined the evolution of genomic expansion in clonal multicellular organisms, using empirically-determined mutational biases for prokaryotes and eukaryotes (12). We modeled a scenario where the fitness benefit of genomic expansion increases with organismal size, both reflecting the fact that larger organisms have more potential ways to divide labor and coordinate morphogenesis, and to examine the scenario where selection very strongly favors an increase in both organismal and genome sizes. Despite increasingly strong selection for genome expansion, larger multicellular organisms were uniformly less capable of evolving larger genomes in our simulation (Figure 1d). This was due entirely to increased genetic drift resulting from a multicellular life history. This decline was more severe for prokaryotes, and at large size, they alone were completely incapable of genomic expansion.

Whether drift-driven genome degradation places an insurmountable constraint on the evolution of complex multicellularity in prokaryotes is yet to be determined. Future work should test, and extend, the predictions of this hypothesis by examining patterns of genomic evolution across independent unicellular-multicellular transitions, modifying *Ne* directly in laboratory evolution experiments, and generating first-principles theoretical predictions via mathematical modeling. Our hypothesis, if supported by further work, stands to rewrite the prevailing narrative for why complex multicellularity repeatedly evolves in eukaryotes yet never in prokaryotes.

## Supporting information

Supplementary data and scripts to procedurally generate all figures

## Materials and Methods

### Cyanobacterial genomic analysis

Prior work by Hammerschmidt et al. (2021) examined the evolution of multicellularity in cyanobacteria. The character trait matrix from Supplementary Table 1 in Hammerschmidt et al. (2021) was used to generate the plots in Figures 1b and c. The plot in Figure 1b is a regression of the sum of the total number of multicellular traits any an individual species identified in Hammerschmidt et al. (2021) contained (out of a maximum of 5: filamentous growth, gas vesicles, true branching, akinetes and hormogonia), against total genome size. 199 genotypes were represented in this dataset, representing all major cyanobacterial phylogenetic groups. Data for total genome size in MB was obtained by querying the accession numbers listed in Supplementary Table 1 against the NCBI Genome Assembly database, taking the genome size from the RefSeq database. We also obtained data from NCBI on the number of pseudogenes (available for 99/199 genomes), and expressed this as the proportion of pseudogenes and protein coding genes in the genome (Figure 1c). All raw data used in this paper is included in the supplement.

### Wright-Fisher simulation of genome size evolution

To examine how broadly the evolution of clonal multicellularity constrains the evolution of larger genome size in prokaryotes and eukaryotes, we developed a simulation based on the Wright-Fisher model, a standard technique in population genetics. Our simulation tracks the genome size of clonal groups of cells over time. Each generation, genotypes are selected to reproduce with fitness-weighted probabilities (*P*). In this simulation, increased genome size is adaptive. We wanted to explore the scenario where selection strongly favors the joint evolution of larger genome size and larger organismal size, as these are universal characteristics of complex multicellularity. We thus included a positive interaction between organismal size and genome size: larger organisms benefit more from having larger genomes, reflecting the potential for there being more ways to divide labor and coordinate morphogenesis in larger organisms.

Specifically, the probability that a genotype is selected in our simulation, *P*_*i*_, is generated using a simple non-linear function, *F*,

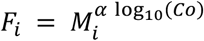

which is normalized such that all selection probabilities sum to 1:

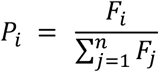

Where *M*_*i*_ is genome size formed by genotype *i, Co* is its group size, and *α* is a scaling factor used to tune the strength of the positive interaction between genome size and group size. In the context of a Wright-Fisher model, this relationship must be nonlinear: because all fitness-weighted probabilities of selection must sum to 1, linear scaling functions have no effect. This function is not meant to capture actual relationships between group size and genome size in nature, but instead is a simple, transparent, and tunable way of exploring the ability for selection to act on increased genome sizes as a function of organismal size.

The genotypes all start the simulation with the same genome size, normalized to 1, and when a multicellular individual reproduces, it has a chance of mutation inserting or deleting a fixed amount (0.1% of the original genome size, roughly the size of a single gene in a typical prokaryotic genome) from its genome. The ratio of deletion to insertion is controlled by a parameter in our simulation called indel_ratio. We used empirically-determined estimates of this ratio from (12): 5 for prokaryotes and 0.8 for eukaryotes.

The total number of cells in the population remains fixed at 10^6^, so the formation of larger groups means the population contains fewer groups in total. We assume a single cell genetic bottleneck during group-level reproduction (*B* = 1, in the context of Equation 1 and Figure 1a). The model was run for 10,000 generations, with 20 replicate simulations for every parameter combination. The mean fold change in genome size (averaged over the groups in each simulation and then over the replicates) is plotted against group size (*Co*) in Figure 1d.

The simulation was written in Python, using numpy. All figures were plotted in Python 3.10, using matplotlib and seaborn. The Large Language Model ChatGPT 4 (OpenAI, 10.25.2023 version), was used to modify Python code for figure plotting via using the “Advanced Data Analysis” plugin. The Python scripts and notebooks used to run the simulation and generate the plots for Figure 1 are available in the supplement. Large Language Model Claude2-100k (Anthropic) was used to proofread the manuscript and make suggestions for technical writing improvements (*i*.*e*., spelling, grammar, and to a limited extent, sentence structure). All ideas in this paper are our own.

## Acknowledgements

We would like to thank Profs. Michael Lynch and Howard Ochman for giving outstanding talks that seemed irreconcilable at first, but stimulated the core ideas behind this paper. Michael Lynch, Peter Conlin, Ozan Bozdag, Kai Tong, Eric Libby and Sam Brown provided stimulating discussions, healthy push-back, and helpful comments on the manuscript. This work was supported by grants from the NIH (Grant No. 5R35GM138030 and T32GM142616) and the NSF Division of Environmental Biology (Grant No. DEB-1845363).

## Description of Supplementary Data

All scripts and data required to procedurally generate Figure 1a-d are supplied in the supplement. This includes 7 files:

1. Cyanobacterial data from Hammerschmidt et al (2021), in a table format used for this paper. In addition, we have added genome size and pseudogene data from NCBI.
2. Scripts to analyze this data, along with the models in Figure 1a-d (four separate files).
3. The outcome of 10,000 generations of the Wright-Fisher model, code to run this model described above.
4. A Jupyter notebook (“Run to Generate All Figures.ipynb”) to run all four models/data analyses and plot them, regenerating Figure 1 procedurally.

