## Supplementary data and scripts to procedurally generate all figures for "A non-adaptive explanation for macroevolutionary patterns in the evolution of complex multicellularity": 1a.py

```
# -*- coding: utf-8 -*-
"""
Created on Wed Oct 11 01:32:45 2023

@author: wratcliff3
"""

import numpy as np
import matplotlib.pyplot as plt

# Define the range of organism sizes (Co)
Co = np.linspace(1, 10000, 100)

# Define different bottleneck sizes (B)
Bs = [1, 10, 100, 500, 1000]

plt.figure(figsize=[8, 8])
for B in Bs:
    proportion = np.minimum(B, Co) / Co
    reciprocal_proportion = 1 / proportion  # Calculate the reciprocal of the proportion
    plt.plot(Co, reciprocal_proportion, label=f'B = {B}')  # Plot the reciprocal proportion

plt.xlabel('Organism Size (Co)', fontsize=24)
plt.ylabel('Fold reduction in Max Diversity (N/L)', fontsize=24)  # Updated ylabel
plt.yscale('log')  # Setting y-axis to logarithmic scale

# Increasing the size of the x and y axis numerical labels to 16 point.
plt.xticks(fontsize=16)
plt.yticks(fontsize=16)

# Adjust the legend size to 16 point.
plt.legend(bbox_to_anchor=(0., 1.02, 1., .102), loc='lower left', ncol=len(Bs), mode="expand", 
           borderaxespad=0., handlelength=1, fontsize=16)

plt.grid(False)
plt.tight_layout()
plt.savefig("Fig1a.pdf")
plt.show()
```
