## Supplementary data and scripts to procedurally generate all figures for "A non-adaptive explanation for macroevolutionary patterns in the evolution of complex multicellularity": 1b.py

```
# -*- coding: utf-8 -*-
"""
Created on Wed Oct 11 01:39:25 2023

@author: wratcliff3
"""

import pandas as pd
import matplotlib.pyplot as plt
import seaborn as sns
import statsmodels.api as sm

# Read the data
data = pd.read_csv("Cyano_Data.csv")

# Convert "Genome size" to megabases (MB)
data["Genome size (MB)"] = data["Genome size"] / 1e6

# Plot "Genome Size (MB)" against "Number MC traits" with linear regression
plt.figure(figsize=[8, 8])
sns.regplot(x=data["Number MC traits"], y=data["Genome size (MB)"], scatter_kws={'s':5, 'label':'Data Points'}, line_kws={'color':'blue'})

# Styling
plt.ylabel('Genome Size (MB)', fontsize=24)
plt.xlabel('Number of Multicellular Adaptations', fontsize=24)
#plt.legend(bbox_to_anchor=(0., 1.02, 1., .102), loc='lower left', mode="expand", borderaxespad=0.)
plt.grid(False)
plt.xticks(fontsize=16)
plt.yticks(fontsize=16)

# Save the plot as a PDF file
plt.tight_layout()
plt.savefig("Fig1b.pdf")

# Display the plot
plt.show()

# Perform linear regression analysis using the converted Genome size in MB
X = sm.add_constant(data["Number MC traits"])  # adding a constant
model = sm.OLS(data["Genome size (MB)"], X).fit()
print(model.summary())
```
