## Supplementary data and scripts to procedurally generate all figures for "A non-adaptive explanation for macroevolutionary patterns in the evolution of complex multicellularity": 1c.py

```
# -*- coding: utf-8 -*-
"""
Created on Wed Oct 11 01:53:51 2023

@author: wratcliff3
"""

import pandas as pd
import matplotlib.pyplot as plt
import seaborn as sns

# Read the data
data = pd.read_csv("Cyano_Data.csv")

# Set the size and style of the plot
plt.figure(figsize=[3, 8])
sns.set_style("whitegrid")

# Create the violin plot with lighter palette and contrasting points
sns.violinplot(x=data["11 FILAMENTOUS"], y=data["Proportion Pseudogenes"], inner="point", 
               palette="Blues_r",  # Using a reversed blue palette for lighter shades
               scale="width", cut=0, 
               linewidth=0.5,  # Thin boundary for the violins
               scatter_kws={"color": "red", "s": 15})  # Contrasting red for the points

# Set y-axis limit and labels for x-axis
plt.ylim(bottom=0)
plt.xticks([0, 1], ["Unicellular", "Multicellular"])
plt.xlabel('')  # Removing the default x-axis label
plt.ylabel('Proportion Pseudogenes', fontsize=24)
plt.xticks(fontsize=16)
plt.yticks(fontsize=16)
# Angle the x-axis labels
plt.xticks(rotation=45, ha='right')
plt.tight_layout()
plt.savefig("Fig1c.pdf", bbox_inches='tight')

plt.show()
```
