## Supplementary data and scripts to procedurally generate all figures for "A non-adaptive explanation for macroevolutionary patterns in the evolution of complex multicellularity": 1d.ipynb

```
# %% [markdown]
By: Emma Bingham

Date: Oct. 27, 2023

# %% [code]
import numpy as np
import matplotlib.pyplot as plt
from tqdm.notebook import tqdm
from matplotlib.lines import Line2D
import json

# %% [code]
def genome_fitness(genomes, group_size, factor=1):
    """ 
    Calculate fitness of genomes and produce probabilities for selection.
    
    Want to nonlinearize the probabilities based on group size and another factor.
    They start off as linear, and we want to bend them to be weighted higher at the top of the distribution
       and lower at the bottom.
       
    factor: the factor B that tunes the strength of the selection
    """
    
    probs = genomes/genomes.sum() # normalize to produce initial linear probabilities
    
    new_probs = probs ** (factor * np.log10(group_size)) # apply equation for nonlinearizing

    return new_probs/new_probs.sum() # normalize again to produce final probabilities

# %% [markdown]
Visualization of the effect of group size and B on the selection probabilities.

# %% [code]
fig, axes = plt.subplots(1,3,figsize=(15,5), dpi=100)

test_genomes = np.random.normal(loc=1, scale=0.1, size=20) 

for j, B in enumerate([0.2, 4, 8]):
    ax = axes[j]
    ax.plot(test_genomes, test_genomes/test_genomes.sum(), color="k")
    ax.set_title(f"B = {B}")
    for num in [10, 100, 1000]:
        test_probs = genome_fitness(test_genomes, num, factor=B)
        ax.scatter(test_genomes, test_probs, label=f"Group size = {num}") 
        ax.axhline(0, linestyle="--", c="k")
        ax.set_xlabel("Genome size")
        ax.set_ylabel("Probability distribution")
        ax.legend()
        ax.set_ylim(-0.1, 0.5)
fig.tight_layout()

# %% [code]
def wright_fisher_with_selection(pop_size=100000, num_groups=10000, initial_genome_size=1, 
                           num_gens=1000, genome_increment=0.001, mu=0.01, indel_ratio=5, factor=1):
    """
    pop_size: population size
    num_groups: number of groups (must be smaller than pop_size)
    initial_genome_size: how big of a genome each group should start with
    num_gens: number of generations to run the Wright-Fisher simulation
    genome_increment: how much to increase or decrease the genome
    mu: mutation rate (e.g. 1 in 100), probability to gain or lose a genome increment
    indel_ratio: ratio of deletions to insertions
    factor: also called B. How strongly selection will select larger genomes
    """
    
    group_size = int(pop_size/num_groups)
    
    # probabilities for [same, increased, decreased] genome
    genome_probs = [1-(mu*(1+indel_ratio)), mu, indel_ratio*mu]    
            
    # groups are represented by genome_size
    groups = initial_genome_size * np.ones(shape=num_groups, dtype=np.float64)
    
    population_history = [groups]
    
    for i in range(num_gens):
                
        # produce probabilities for selection
        probs = genome_fitness(groups, group_size, factor=factor)
        
        # select groups for next generation
        groups = groups[np.random.choice(num_groups, replace=True, p=probs, size=num_groups)]
        
        # mutate genomes
        # there's a chance of getting either bigger or smaller
        # and a larger chance of staying the same
        # biased towards smaller for prokaryotes, bigger for eukaryotes
        groups = groups + np.random.choice([0, genome_increment, -genome_increment], 
                                       size=num_groups, 
                                       p=genome_probs)

        
        population_history.append(groups)
        
    return np.array(population_history)

# %% [code]
parameters = {
    "num_gens": 1000, #10000,
    "pop_size": int(10**6),
    "initial_genome_size": 1,
    "genome_increment": 0.001, # amount of genome to add when a group mutates (+ for eukaryotes, - for prokaryotes)
    "mu": 0.01, # probability to increase or decrease genome
}

pro_indel_ratio = 5 # deletion/insertion, inspired by Ochman paper (Kuo and Ochman 2009), between 5 and 10 for prokaryotes
eu_indel_ratio = 0.8 # inspired by Ochman paper (Kuo and Ochman 2009)

reps = 20
Bs = [2, 4, 8]

# %% [code]
nums_groups = np.array([10, 20, 50, 100, 200, 500, 1000, 2000, 5000, 10000, 20000, 50000, 100000])
print(nums_groups)

results = {fac:[] for fac in Bs}

for fac in Bs:
    parameters["factor"] = fac
    pro = {}
    eu = {}
    for num_groups in tqdm(nums_groups):
        parameters["num_groups"] = num_groups

        pro_reps = []
        eu_reps = []
        for rep in range(reps): # run reps number of simulations
            
            # just save the final genome values
            parameters["indel_ratio"] = pro_indel_ratio
            pro_reps.append(wright_fisher_with_selection(**parameters)[-1,:])

            parameters["indel_ratio"] = eu_indel_ratio
            eu_reps.append(wright_fisher_with_selection(**parameters)[-1,:])

        pro[num_groups] = np.array(pro_reps)
        eu[num_groups] = np.array(eu_reps)
    results[fac].append(pro)
    results[fac].append(eu)

# %% [code]
fig, ax = plt.subplots(1,1,figsize=(6,6), dpi=100)

group_sizes = parameters["pop_size"]/np.array(nums_groups)
linestyles = ["-", "--", "dotted"]

for j, fac in enumerate(Bs):
    pro, eu = results[fac]
    pro_final_genomes = [pro[g].mean(axis=1) for g in nums_groups] # mean final genomes of each rep
    eu_final_genomes = [eu[g].mean(axis=1) for g in nums_groups]
    ax.errorbar(group_sizes, [arr.mean() for arr in pro_final_genomes], 
                yerr=[arr.std()/(len(arr)**0.5) for arr in pro_final_genomes], 
                marker="o", c="#e63946", capsize=2, linestyle=linestyles[j])
    ax.errorbar(group_sizes, [arr.mean() for arr in eu_final_genomes], 
                yerr=[arr.std()/(len(arr)**0.5) for arr in eu_final_genomes], 
                marker="o", c="#3a86ff", capsize=2, linestyle=linestyles[j])

ax.set_xlabel("Group Size")
ax.set_ylabel("Fold Change in Genome Size")
ax.semilogx()

custom_lines = [Line2D([0], [0], color="k", linestyle=line, lw=2) for line in linestyles]

ax.legend(custom_lines, [f"B = {fac}" for fac in Bs])

ax.set_title(f"Fold Change in Genome Size After {parameters['num_gens']} generations")

# %% [code]
results_to_json = {int(B): [{int(g): arr.tolist() for g, arr in pro_eu.items()} for pro_eu in res] for B, res in results.items()}
del parameters["num_groups"]

simulation = {
    "parameters": parameters,
    "num_reps": int(reps),
    "Bs": Bs,
    "nums_groups": nums_groups.tolist(),
    "results": results_to_json,
}

# %% [code]
with open("results_500gens.json", "w") as f:
    json.dump(simulation, f) 

# %% [code]
```
