## Supplementary data and scripts to procedurally generate all figures for "A non-adaptive explanation for macroevolutionary patterns in the evolution of complex multicellularity": Run to Generate All Figures.ipynb

```
# %% [code]
import pandas as pd
import numpy as np
import matplotlib.pyplot as plt
from matplotlib.lines import Line2D
import seaborn as sns
import statsmodels.api as sm
import json

# %% [code]
from matplotlib.gridspec import GridSpec

# %% [markdown]
Fig 1a data

# %% [code]
# Define the range of organism sizes (Co)
Co = np.linspace(1, 10000, 100)

# Define different bottleneck sizes (B)
Bs = [1, 10, 100, 500, 1000]

# %% [markdown]
Fig 1b and 1c data

# %% [code]
# Read the data
data = pd.read_csv("Cyano_Data.csv")

# Convert "Genome size" to megabases (MB)
data["Genome size (MB)"] = data["Genome size"] / 1e6

# %% [markdown]
Fig 1d data

# %% [code]
with open("../Figure 1d/results_10000gens.json", "r") as f:
    simulation = json.load(f)

nums_groups = simulation["nums_groups"]
parameters = simulation["parameters"]
factors = simulation["Bs"]
results = {int(B): [{int(g): np.array(arr) for g, arr in pro_eu.items()} for pro_eu in res] for B, res in simulation["results"].items()}

# %% [markdown]
Plotting

# %% [code]
fig = plt.figure(figsize=(10, 10), dpi=150)
# gs = GridSpec(1, 4, width_ratios=[3,2,1,3])
gs = GridSpec(2, 4)

ax1 = fig.add_subplot(gs[0,0:2])
ax2 = fig.add_subplot(gs[0,2:])
ax3 = fig.add_subplot(gs[1,0:1])
ax4 = fig.add_subplot(gs[1,1:])

# Fig 1a
for B in Bs:
    proportion = np.minimum(B, Co) / Co
    reciprocal_proportion = 1 / proportion  # Calculate the reciprocal of the proportion
    ax1.plot(Co, reciprocal_proportion, label=f'B = {B}')  # Plot the reciprocal proportion
    ax1.text(10000, max(reciprocal_proportion), f'B = {B}', fontsize=14, va="bottom", ha="right")

ax1.set_xlabel('Organism Size (Co)', fontsize=16)
ax1.set_ylabel('Fold reduction in Max Diversity (N/L)', fontsize=16)  # Updated ylabel
ax1.semilogy()  # Setting y-axis to logarithmic scale
# ax1.set_xlim(-200, 12000)

# Increasing the size of the x and y axis numerical labels to 16 point.
ax1.tick_params(axis='both', labelsize=12)

# Adjust the legend size to 16 point.
# Decided to remove the legend in favor of on-plot annotations
# ax1.legend(bbox_to_anchor=(0., 1.02, 1., .102), loc='lower left', ncol=len(Bs), mode="expand", 
#            borderaxespad=0., handlelength=1, fontsize=14)
# ax1.legend(bbox_to_anchor=(0., 1.02, 1., .102), ncol=3, mode="expand", 
#            borderaxespad=0., handlelength=1, fontsize=14)
# ax1.legend(bbox_to_anchor=(1.1, 1.2), ncol=1, 
#            borderaxespad=0., handlelength=1, fontsize=14)


ax1.grid(False)


# Fig 1b
# Plot "Genome Size (MB)" against "Number MC traits" with linear regression
sns.regplot(x=data["Number MC traits"], y=data["Genome size (MB)"], 
            scatter_kws={'s':5, 'label':'Data Points'}, line_kws={'color':'blue'}, ax=ax2)

# Styling
ax2.set_ylabel('Genome Size (MB)', fontsize=16)
ax2.set_xlabel('Number of Multicellular Adaptations', fontsize=16)
ax2.tick_params(axis='both', labelsize=12)
ax2.grid(False)
ax2.set_xlim(-0.2, 5.2)
# ax2.set_xticklabels([str(i) for i in range(6)])


# Fig 1c
sns.set_style("whitegrid")

# Create the violin plot with lighter palette and contrasting points
sns.violinplot(x=data["11 FILAMENTOUS"], y=data["Proportion Pseudogenes"], inner="point", 
               palette="Blues_r",  # Using a reversed blue palette for lighter shades
               scale="width", cut=0, 
               linewidth=0.5,  # Thin boundary for the violins
               scatter_kws={"color": "red", "s": 15}, ax=ax3)  # Contrasting red for the points

# Set y-axis limit and labels for x-axis
ax3.set_ylim(bottom=0)
ax3.set_xticklabels(labels=["Unicellular", "Multicellular"], rotation=45, ha="right", fontsize=14)
ax3.set_xlabel('')  # Removing the default x-axis label
ax3.set_ylabel('Proportion Pseudogenes', fontsize=16)

# Make y-axis tick labels larger
ax3.tick_params(axis='y', labelsize=12)


# Fig 1d
group_sizes = parameters["pop_size"]/np.array(nums_groups)
linestyles = ["-", "--", "dotted"]

for j, fac in enumerate(factors):
    pro, eu = results[fac]
    pro_final_genomes = [pro[g].mean(axis=1) for g in nums_groups] # mean final genomes of each rep
    eu_final_genomes = [eu[g].mean(axis=1) for g in nums_groups]
    ax4.errorbar(group_sizes, [arr.mean() for arr in pro_final_genomes], 
                yerr=[arr.std()/(len(arr)**0.5) for arr in pro_final_genomes], 
                marker="o", c="#3a86ff", capsize=2, linestyle=linestyles[j])
    ax4.errorbar(group_sizes, [arr.mean() for arr in eu_final_genomes], 
                yerr=[arr.std()/(len(arr)**0.5) for arr in eu_final_genomes], 
                marker="o", c="#e63946", capsize=2, linestyle=linestyles[j])

ax4.set_xlabel("Organism Size (Co)", fontsize=16)
ax4.set_ylabel("Mean Fold Change in Genome Size", fontsize=16)
ax4.tick_params(axis='both', labelsize=12)
ax4.semilogx()

# make horizontal line at 1
ax4.axhline(1, color="gray", linestyle="-", alpha=0.5)

# make custom legend
custom_lines = [Line2D([0], [0], color="k", linestyle=line, lw=1) for line in linestyles]
ax4.legend(custom_lines, [r"$\alpha$"+f" = {fac}" for fac in factors], loc="lower left", fontsize=14)

# annotate prokaryotes and eukaryotes
ax4.text(10**4, 1.35, "Eukaryotes", fontsize=14, color="#e63946")
ax4.text(4*10**2, 0.9, "Prokaryotes", fontsize=14, color="#3a86ff")

ax4.grid(False)

# put figure (a), (b), (c), (d) labels
ax1.text(-400, 10**4-100, '(a)', fontsize=12, ha='left', va='bottom') #, transform=plt.gca().transAxes);
ax2.text(-0.15, 13.05, '(b)', fontsize=12, ha='left', va='bottom')
ax3.text(-0.23, 0.317, '(c)', fontsize=12, ha='right', va='bottom')
ax4.text(10**5+7000, 1.51, '(d)', fontsize=12, ha='left', va='bottom')

fig.tight_layout()

plt.savefig("Figure_square_final.pdf", dpi=150)

# %% [code]


# %% [code]
```
