## Supplementary figures and images for "A non-adaptive explanation for macroevolutionary patterns in the evolution of complex multicellularity"

### Figure_square_final.pdf

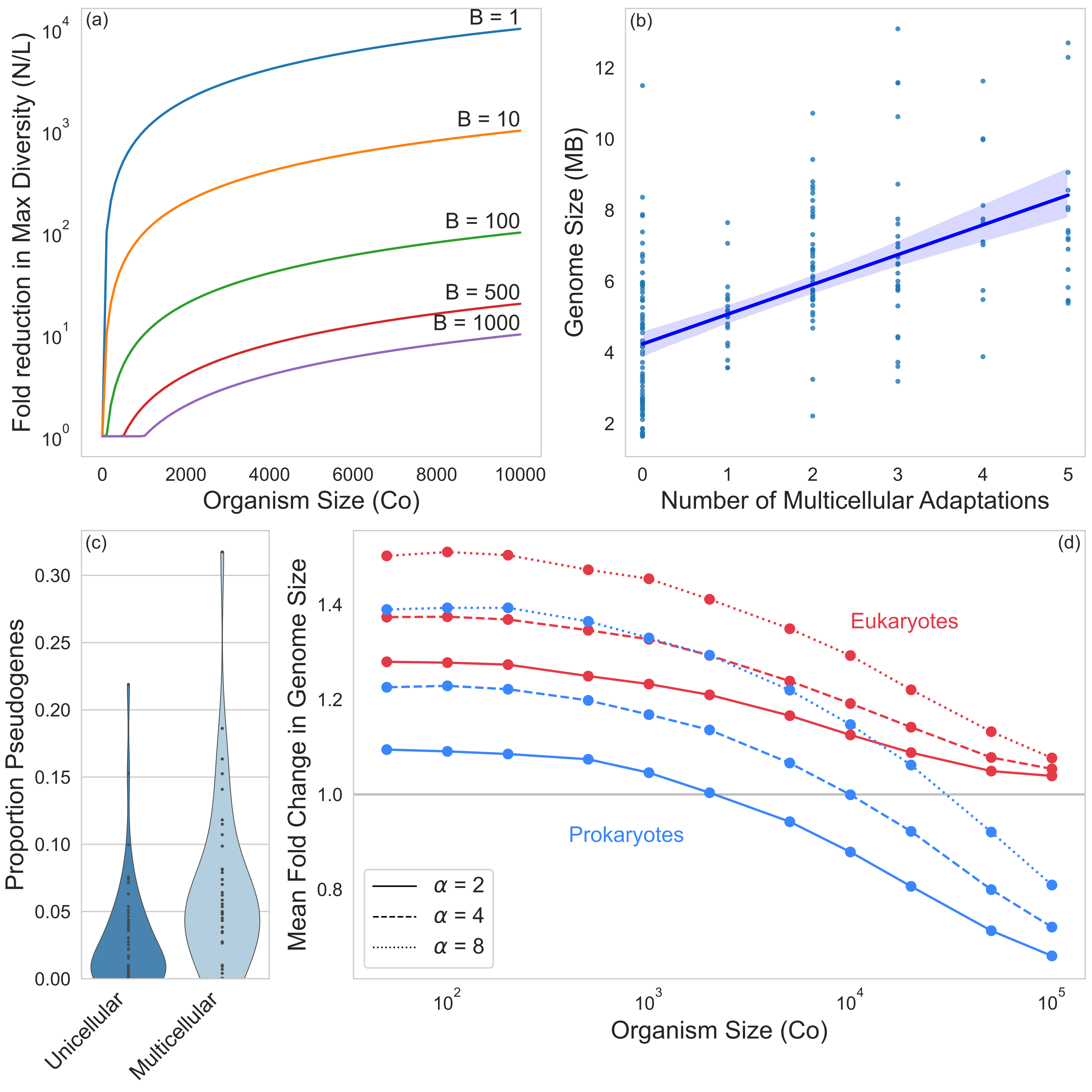
